# Chelerythrine as an anti-Zika virus agent: therapeutic potential and mode of action

**DOI:** 10.1101/2022.12.16.520682

**Authors:** Erica Españo, Jiyeon Kim, Chong-Kil Lee, Robert G. Webster, Richard J. Webby, Jeong-Ki Kim

## Abstract

Zika virus (ZIKV) is a mosquito-borne virus that has been associated with adult and neonatal neurological conditions. So far, there is no approved drug or vaccine against ZIKV infection; thus, ZIKV remains a global health threat. Here, we explored the effects of chelerythrine (CTC), a known protein kinase C (PKC) inhibitor, against ZIKV infection in cell culture models to determine its potential as a therapeutic agent for ZIKV infection. We found that CTC protected Vero cells from ZIKV-induced cytopathic effects in a dose-dependent manner. It also reduced the production of ZIKV in Vero and A549 cells. In contrast, other PKC inhibitors failed to protect Vero cells from ZIKV-induced cytopathic effects, indicating PKC-independent mechanisms underlying the effects of CTC on ZIKV. Further investigation suggested that CTC inhibited ZIKV attachment/binding rather than internalization in the host cells. Pretreatment of cell-free ZIKV particles rather than pretreatment of cells with CTC resulted in reduced ZIKV infectivity *in vitro*, indicating that CTC blocked the attachment/binding of the ZIKV particles to host factors. *In silico* analyses suggested that these effects are potentially due to the binding of CTC to the ZIKV E protein, which may occlude the interaction of the E protein with attachment factors or receptors on the host cell surface. Overall, our findings suggest that CTC reduces the infectivity of ZIKV particles through PKC- and cell-independent mechanisms. Our findings also support further exploration of CTC as an anti-ZIKV agent.

## Introduction

Zika virus (ZIKV) is a mosquito-borne member of the *Flaviviridae* family of positive, single-stranded, enveloped RNA viruses. It is related to other arthropod-borne human pathogens, such as the dengue virus (DENV), Japanese encephalitis virus, West Nile virus (WNV), and yellow fever virus. Most cases of ZIKV infection are asymptomatic (50–80%), and majority of symptomatic cases manifest as mild and self-limiting febrile illness (1,2). However, the large outbreak in Brazil in 2015 revealed associations between ZIKV infection and adult (i.e., Guillain-Barré syndrome) and neonatal (i.e., microcephaly) neurological conditions (3,4,5), leading to the declaration of a Public Health Emergency of International Concern by the World Health Organization (WHO) in February 2016 (6). Later studies revealed that ZIKV infection in expectant mothers can cause a broad spectrum of neurological disorders in addition to microcephaly, which are now collectively called the congenital Zika syndrome (5).

Until the ZIKV outbreak on Yap Island, Federated States of Micronesia in 2004, ZIKV outbreaks were small and mostly confined to Asia and Africa (1). Since the 2015 ZIKV outbreak in South America, local transmission of ZIKV has been reported in 87 countries, although recent clusters have been small (7). The circulation of ZIKV is currently limited to tropical and subtropical regions because of the geographic distribution of *Aedes (Ae*.*) aegypti* mosquitoes, which are considered the primary vectors for urban transmission of ZIKV (8). In addition, *Ae. albopictus* mosquitoes, which inhabit subtropical and temperate regions, have been reported to transmit ZIKV in experimental settings, suggesting that these mosquitoes may contribute to the urban cycle of ZIKV (9,10,11). Although the role of *Ae. albopictus* in the overall spread of ZIKV remains unclear, it is considered a potential vector for spreading ZIKV to subtropical and other regions where human populations are immunologically naïve to ZIKV. The steadily increasing global temperatures are also expected to broaden the geographic ranges of *Ae. aegypti* and *Ae. albopictus*, which may lead to the expansion of locations with autochthonous ZIKV transmission (12).

Given the threat of future ZIKV outbreaks to immunologically naïve populations, WHO considers ZIKV a priority pathogen for research (13). However, to date, there are no approved drugs or vaccines against ZIKV. Academic institutions and pharmaceutical companies alike have turned to drug repositioning for the rapid discovery of anti-ZIKV agents. This strategy involves screening libraries of approved and investigational compounds for therapeutic potential against ZIKV infection to bypass some of the steps in the lengthy and costly drug discovery and development pipeline. We have previously reported a primary screening of over 2,000 approved and investigational compounds that led to the identification of lipophilic statins as inhibitors of ZIKV infection *in vitro (*14). Chelerythrine (in the form of chelerythrine chloride; CTC), a cell-permeable benzo[c]phenanthridine alkaloid under investigation as an anticancer agent owing to its ability to inhibit protein kinase C (PKC), also presented as a candidate on the same primary screen. Here, we performed a series of infectivity assays on Vero and A549 cells to demonstrate the anti-ZIKV effects of CTC *in vitro* and to identify potential mechanisms underlying its effects. The findings presented here suggest that CTC should be considered as a potential candidate for the treatment of ZIKV infection.

## Results

### CTC inhibits ZIKV infection in vitro

Primary phenotypic screening of over 2,000 compounds revealed the anti-ZIKV activity of CTC in Vero cells. To validate the results of the primary screening, we determined the effects of different concentrations of CTC on ZIKV infection at a multiplicity of infection (MOI) of 0.02 in Vero cells using a colorimetric cytotoxicity assay. CTC was added at the same time as ZIKV, and neither the drug nor the virus was removed during the five-day incubation period to ensure that CTC could act on all potential target stages in the ZIKV replication cycle. CTC protected Vero cells from ZIKV-induced cytopathic effects (CPE) in a dose-dependent manner (**Fig. 1A**) at non-cytotoxic concentrations (**Fig. 1B**). The calculated half-maximal effective concentration (EC_50_) of CTC against ZIKV-induced CPE in Vero cells was 692.4 nM (**Fig. 1C**), which coincides with the reported half-maximal inhibitory concentration of CTC for PKC (700 nM) (15). The calculated half-maximal cytotoxic concentration (CC_50_) of CTC in Vero cells was 4.15 μM (**Fig. 1C**), indicating a selectivity index of 6.0 in this system.

**Figure 1.**
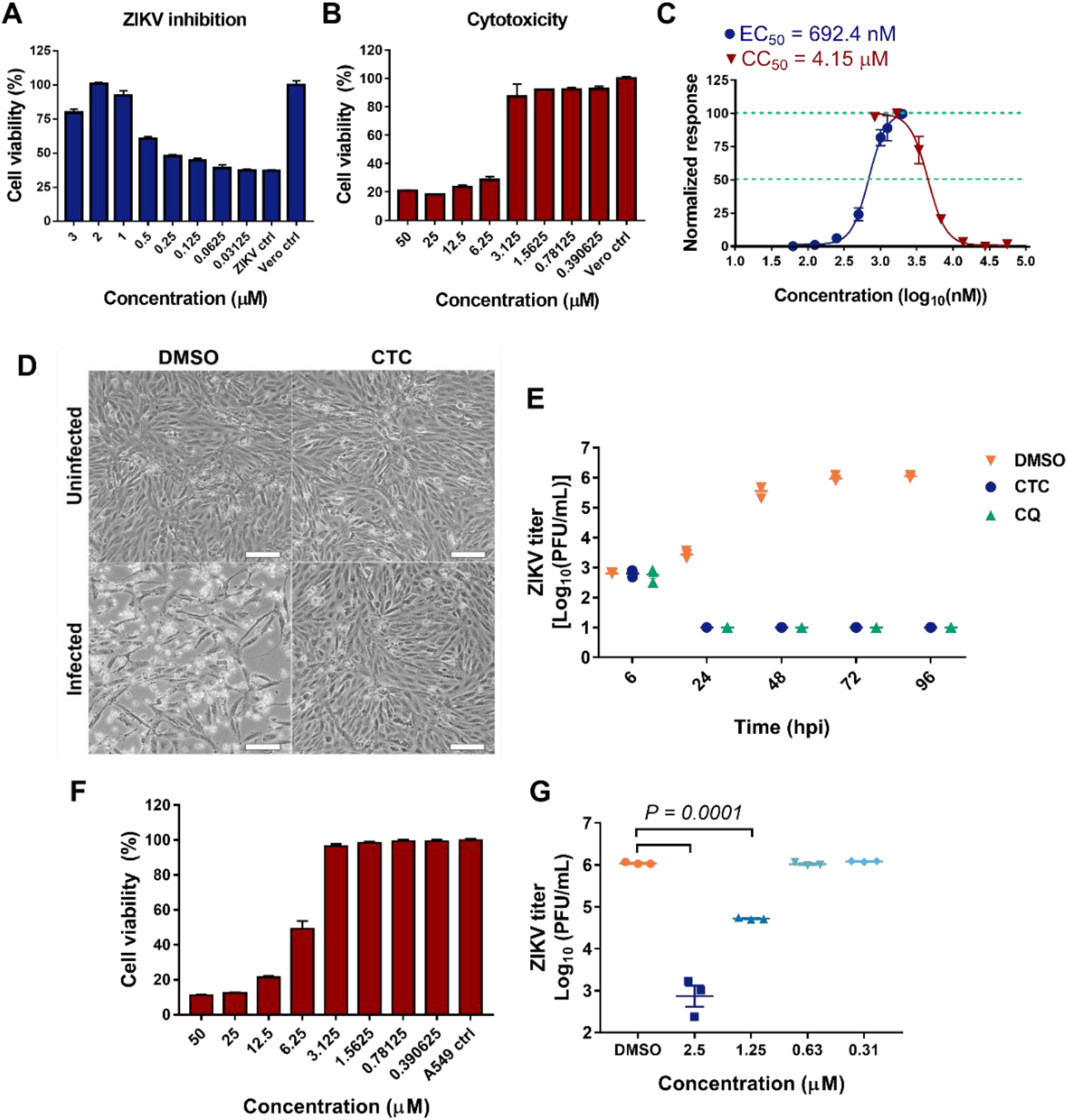
Chelerythrine (CTC) inhibits Zika virus (ZIKV) infection in Vero and A549 cells. Effects of different CTC concentrations on **(A)** cytopathic effects induced by ZIKV infection at a multiplicity of infection (MOI) of 0.01 in Vero cells and **(B)** Vero cell viability. CTC was added at the same time as ZIKV and was not removed during the entire incubation period. Infected Vero cells treated with 1% dimethyl sulfoxide (DMSO) were used as control (ZIKV ctrl). Cell viability was calculated relative to the uninfected Vero cells treated with 1% DMSO (Vero ctrl). Data presented are representative of at least 3 independent experiments. **(C)** Relative half-maximal effective (EC_50_) and cytotoxic (CC_50_) concentrations of CTC against ZIKV-induced cytopathic effects (CPE) and Vero cell viability, respectively. Datapoints are means of normalized data from three independent assays ± standard error of the mean (SEM). **(D)** Effects of 1.45 μM CTC and 0.3% DMSO on Vero cell morphology (Uninfected) and CPE induced by ZIKV infection (Infected, MOI of 0.01). Scale bar: 1000 μm. **(E)** Effects of 1.45 μM CTC, 25 μM chloroquine (CQ), or 0.3% DMSO on ZIKV production in Vero cells over time (MOI of 0.01). **(F)** Effects of CTC on A549 cell viability. Values were calculated relative to cells treated with 1% DMSO (A549 ctrl). Results presented are representative of at least 3 independent experiments. **(G)** Effects of different concentrations of CTC on ZIKV production in A549 cells. CTC was added at the same time as ZIKV (MOI of 0.01) and was not removed during the entire incubation period. All datapoints represent the mean ± SEM; *N =* 3 per concentration. *P-*values indicate significant differences; analysis of variance (ANOVA) followed by Tukey’s multiple comparison test.

Next, we determined the CTC concentration for use in subsequent assays. Treatment with 1.45 μM CTC protected Vero cells from ZIKV-induced CPE without causing apparent changes to cell morphology (**Fig. 1D**). We used this concentration to treat ZIKV-infected Vero cells (MOI of 0.01) and monitored the effects of CTC treatment on ZIKV production over time (**Fig. 1E**). We found that CTC treatment reduced ZIKV production to undetectable titers as early as 24 h post-infection (hpi), which was similar to the effects of 25 μM chloroquine (CQ), a reported ZIKV endocytosis inhibitor (14,16). The level of ZIKV reduction by CTC was sustained throughout the incubation period. We also treated A549 cells, another cell line used as a model for ZIKV infection (17), with non-cytotoxic concentrations of CTC (**Fig. 1F**) in the presence of ZIKV (MOI of 0.01). We found that 3-day treatment with CTC reduced ZIKV production in A549 cells in a concentration-dependent manner (**Fig. 1G**). Notably, treatment with 2.5 μM CTC inhibited ZIKV production in A549 cells by approximately 1000-fold. These results indicate that CTC inhibits ZIKV production *in vitro*.

### Other PKC inhibitors do not inhibit ZIKV infection in Vero cells

The anticancer properties of CTC are primarily attributed to its ability to inhibit conventional (α, βI, βII, and γ), novel (δ, θ, ε, and η), and atypical PKCs (ζ and λ/ι) (18,19). Together with CTC, other PKC inhibitors have been reported to inhibit WNV infection *in vitro*, although the specific PKC isoform required for WNV infection was not identified (20). CTC and other broad-spectrum PKC inhibitors have also been reported to inhibit Rift Valley fever virus (RVFV) infection in cell culture models likely through the inhibition of PKCε (21). Therefore, we tested other PKC inhibitors for anti-ZIKV activity in the hope of identifying more compounds that inhibit ZIKV infection. To identify the PKC isoforms relevant to ZIKV replication, we chose PKC inhibitors with different specificities to the PKC isoforms. Like CTC, bisindolylmaleimide I is considered a pan-PKC inhibitor, although bisindolylmaleimide I has been reported to be more selective towards PKCα and PKCβI (22). Calphostin C, rottlerin, and Gö 6983 are also considered broad-spectrum PKC inhibitors, although they exhibit different potencies towards the different isoforms (23,24,25,26). Gö 6976 inhibits conventional PKC isoforms, especially PKCα and PKCβI (27). Safingol (ʟ-*threo*-dihydrosphingosine) inhibits conventional and novel PKC isoforms (28,29).

To determine the potential dose-dependent protective effects of the PKC inhibitors against ZIKV-induced CPE in Vero cells, we chose a wide range of concentrations, including an upper limit where the compounds showed cytotoxic effects. Remarkably, most of the other PKC inhibitors that we tested, including two PKC inhibitors (calphostin C and rottlerin) that were reported to inhibit RVFV and WNV *in vitro*, did not inhibit ZIKV-induced CPE in Vero cells even at the highest tested concentrations (**Fig. 2**). Only safingol, a lyso-sphingolipid analog, protected Vero cells from ZIKV-induced CPE, but only at a modest degree at the highest non-cytotoxic concentration (**Fig. 2F**). These results suggest that PKC inhibition is not a viable strategy for inhibiting ZIKV infection, and that CTC may inhibit ZIKV infection through PKC-independent mechanisms.

**Figure 2.**
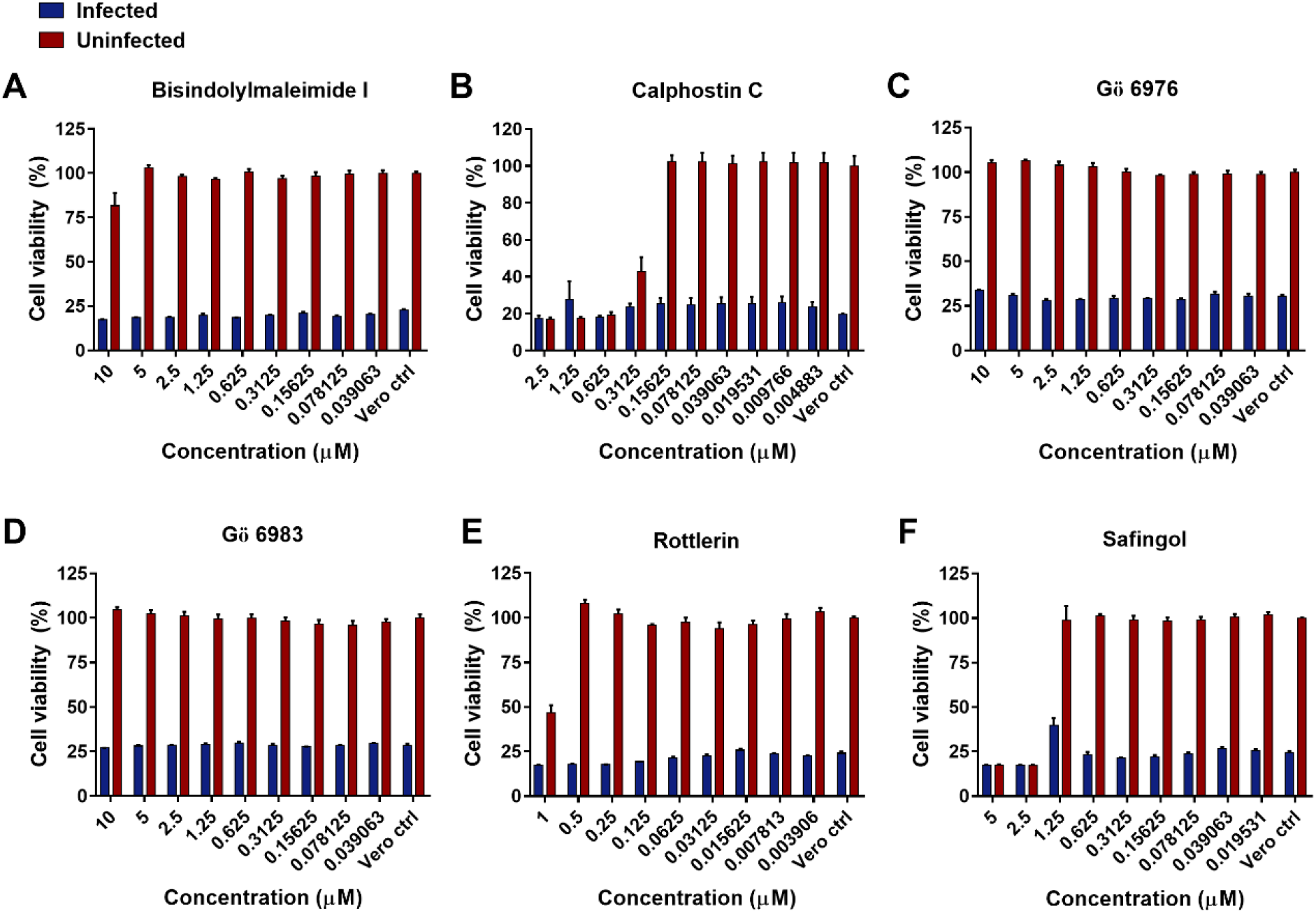
Other protein kinase C (PKC) inhibitors do not inhibit ZIKV infection in Vero cells. Effects of different concentrations of **(A)** bisindolylmaleimide I, **(B)** calphostin C, **(C)** Gö 6976, **(D)** Gö 6983, **(E)** rottlerin, and **(F)** safingol on ZIKV-induced cytopathic effects in Vero cells. Vero cells were infected with MOI of 0.02 of ZIKV and simultaneously treated with PKC inhibitors. Neither the virus nor the PKC inhibitor was removed during the 5-day incubation period. Vero cell controls (Vero ctrl) were treated with 1% DMSO. Cell viability was calculated relative to the uninfected Vero cell controls. Data shown are representative of at least 2 independent experiments. Each datapoint represents the mean ± SEM; *N =* 3 per concentration.

### CTC inhibits ZIKV attachment and/or binding in vitro

To determine the target of CTC in the ZIKV replication cycle, we evaluated the effects of adding CTC at different time points post-infection on a single round of ZIKV replication. Addition of CTC to ZIKV-infected Vero cells (MOI of 10) at 0 to 12 hpi significantly inhibited ZIKV production (**Fig. 3A**). The effects of CTC addition from 0 to 6 hpi were also superior to the effects of CQ on ZIKV production at the corresponding time points. These results suggest that CTC inhibited early rather than late stages of the ZIKV replication cycle.

**Figure 3.**
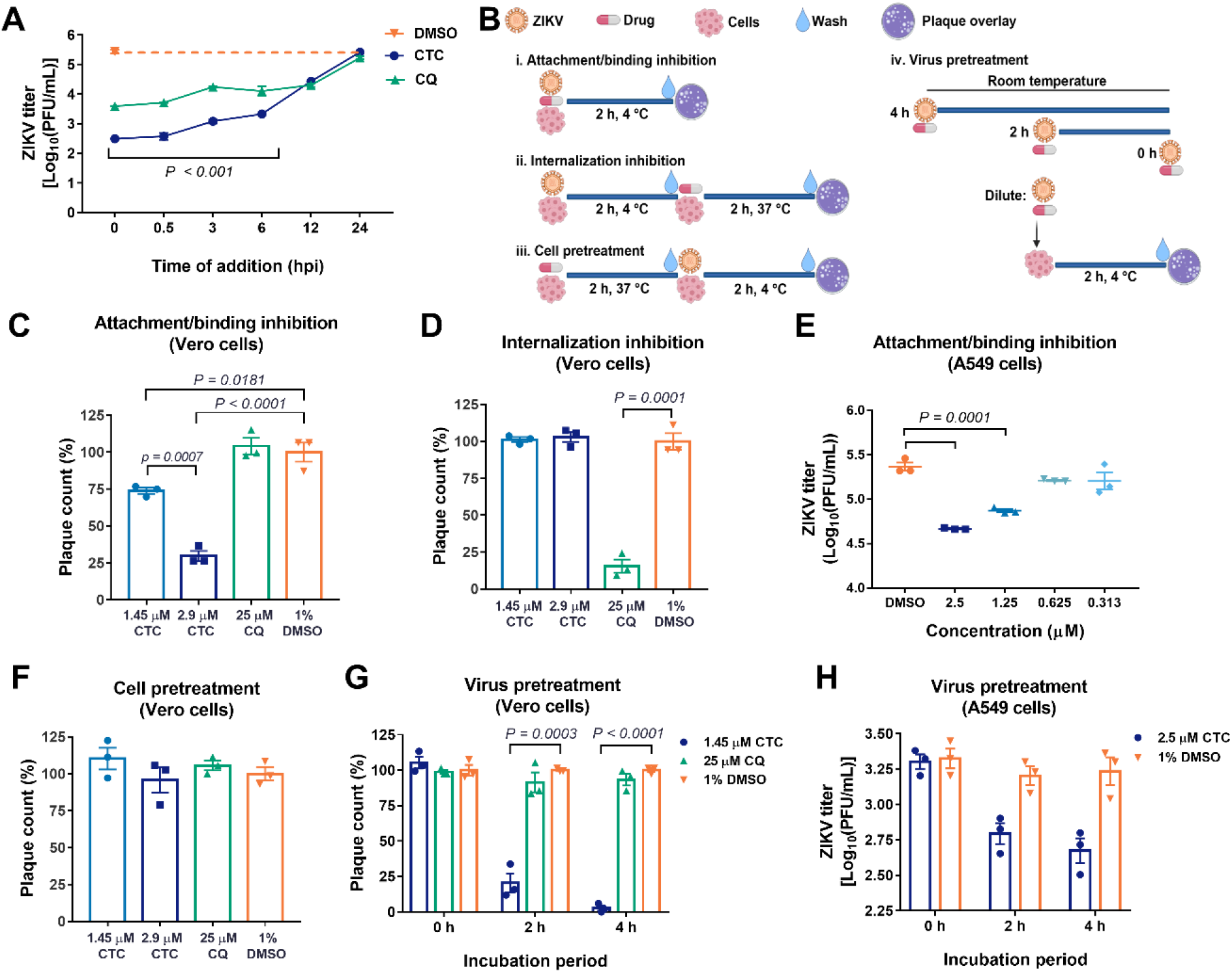
CTC reduces the infectivity of ZIKV particles and inhibits ZIKV binding/attachment to Vero and A549 cells. **(A)** Effects of the addition of 1.45 μM CTC, 25 μM CQ, or 0.3% DMSO at different timepoints post-infection with ZIKV (at a multiplicity of infection of 10) on ZIKV production in Vero cells. Datapoints represent the mean ± SEM; *N* = 3 per timepoint. Hpi, hours post-infection. **(B)** Schematic diagram for the assays performed to determine the targets of CTC on early stages of ZIKV infection in Vero cells. Assays i and iv were similarly performed on A549 cells, except that the A549 cells were incubated without plaque overlay. Instead, the A549 culture supernatants were collected at 24 hpi, and ZIKV production in A549 cells was determined via plaque assay in Vero cells. **(C,D)** ZIKV plaque formation in Vero cells following treatment with CTC, CQ, or DMSO during the **(C)** attachment/binding and **(D)** internalization stages of infection. Data shown are representative of at least 2 independent experiments. **(E)** ZIKV production in A549 cells following treatment with different concentrations of CTC during the binding/attachment stage of infection. **(F,G)** ZIKV plaque formation in Vero cells following pretreatment of **(F)** cells and **(G)** ZIKV particles with CTC, CQ, or DMSO. Data shown are representative of at least 2 independent experiments. (**H)** ZIKV production in A549 cells following pretreatment of ZIKV particles with CTC. For C–H, individual datapoints (*N* = 3 per treatment/timepoint) and means ± SEM are shown. Plaque counts (%) were calculated relative to the mean plaque counts of the DMSO controls. All ZIKV titers are presented as the log_10_ of plaque-forming units (PFU) per mL. *P-*values denote significant differences, two-tailed Student’s *t-*test. The schematic diagram (B) was created with BioRender.com.

To identify the target of CTC in the entry stage of the ZIKV infection cycle, we performed a series of ZIKV attachment/binding and internationalization inhibition assays in Vero and A549 cells (**Fig. 3B**). To determine the effects of CTC on ZIKV attachment/binding, we added different concentrations of CTC and ZIKV (400 plaque-forming units, PFU) to Vero cells and incubated the cells at 4 °C to delay and synchronize virus internalization. We found that CTC treatment during the attachment/binding stage of ZIKV infection reduced plaque formation in Vero cells in a concentration-dependent manner, indicating the reduced ability of ZIKV particles to attach and bind to the cells following CTC treatment (**Fig. 3C**). In contrast, CQ treatment during the attachment/binding step did not reduce ZIKV plaque formation in Vero cells. Next, to determine the effects of CTC on ZIKV internalization, we first allowed ZIKV (400 PFU) to attach to Vero cells at 4°C and then removed the unadsorbed virus particles. We then added the drugs to the infected cells and incubated them at 37 °C to allow viral internalization. Virus particles that were not internalized were inactivated by treatment with citric acid (pH 3.0). Addition of CTC at this stage of infection did not reduce ZIKV plaque formation, which suggests that CTC does not inhibit ZIKV internalization in Vero cells. Meanwhile, CQ, a reported ZIKV endocytosis inhibitor, significantly inhibited ZIKV internalization, as indicated by reduced plaque formation (**Fig. 3D**). These results suggest that CTC can inhibit ZIKV attachment and/or binding to, but not internalization in, Vero cells.

We also tested the effects of CTC on ZIKV attachment and binding to A549 cells. The cells were infected with ZIKV (MOI of 1.0) in the presence of varying concentrations of CTC for 2 h at 4 °C (**Fig. 3B**). Culture supernatants were collected at 24 hpi and titrated through the plaque assay in Vero cells. Similarly, CTC treatment during the ZIKV attachment/binding stage reduced ZIKV production in A549 cells in a concentration-dependent manner (**Fig. 3E**). Altogether, these findings indicate that CTC inhibits ZIKV attachment/binding to host cells.

### CTC reduces the infectivity of ZIKV particles

Because our results using several PKC inhibitors suggest PKC-independent mechanisms underlying the anti-ZIKV activity of CTC, we speculated that cell-independent mechanisms are responsible for the inhibitory effects of CTC on ZIKV attachment/binding. Confirming this, 2-h pretreatment (37 °C) of Vero cells with different concentrations of CTC did not affect ZIKV attachment/binding to Vero cells (**Fig. 3B,F**). To determine whether CTC targets the ZIKV particles rather than the cells, we incubated ZIKV (3.75 × 10^4^ PFU) with 1.45 μM CTC, 25 μM CQ or 1% dimethyl sulfoxide (DMSO) in the absence of cells for 0, 2, and 4 h at room temperature (**Fig. 3B**). The compound + ZIKV mixtures were diluted 1:100 to reduce CTC concentration to subtherapeutic levels (0.015 μM) immediately before allowing the pretreated ZIKV particles to adsorb to Vero cells (2 h, 4 °C). No loss of ZIKV infectivity was observed over the 4-h incubation period, as the mean plaque count of the DMSO group at each time point remained at approximately 50 PFU (**Fig. 3G**). Notably, we observed that longer incubation of ZIKV particles with CTC (4 h) led to lower ZIKV plaque counts. In contrast, pretreatment of ZIKV particles with CQ did not reduce ZIKV attachment/binding to Vero cells. The subtherapeutic levels of CTC without prior incubation with ZIKV (0-h incubation) also did not inhibit ZIKV attachment/binding to the cells, indicating that the reduction in ZIKV infectivity was caused by ZIKV pretreatment with CTC rather than by residual effects of CTC on the cells.

We also tested whether pre-incubating ZIKV particles with CTC affected ZIKV attachment/binding to A549 cells. The ZIKV particles (8 × 10^4^ PFU) were similarly incubated with 2.5 μM CTC for 0, 2, and 4 h prior to adsorption to A549 cells (**Fig. 3*B***). Immediately before adsorption, the CTC + ZIKV mixtures were diluted (1:80) to a sub-therapeutic concentration of CTC (0.031 μM). The culture supernatants of the A549 cells were collected at 24 hpi, and ZIKV titers were determined via plaque assay in Vero cells. Similarly, preincubation of ZIKV with CTC tended to reduce the ability of ZIKV particles to attach and bind to A549 cells (**Fig. 3*H***). Likewise, no loss of ZIKV particles was observed after 4 h of pre-incubation with DMSO. These data suggest that CTC reduces the infectivity of ZIKV particles, leading to the failure of ZIKV particles to attach and bind to host cells.

### CTC may bind to the ZIKV envelope (E) protein

Viral inactivators may reduce the infectivity of enveloped viral particles either by blocking the attachment/binding of the virus by binding envelope glycoproteins or by destroying the viral particles altogether. The ZIKV entry process is highly dependent on the ZIKV E protein, which facilitates attachment to host factors, binding to host receptors, and fusion with the host membrane. Owing to the central role of the E protein in ZIKV attachment and binding to the host cell, we speculated that the ability of CTC to inhibit ZIKV attachment/binding was due to the binding of CTC to the ZIKV E protein. Prediction of the interaction between CTC and the prefusion form of the ZIKV E protein (PDB: 5JHM) (30) suggested that CTC binds a region within the central dimer interface of the ZIKV E protein, with a predicted binding affinity of –7.0 kcal/mol (**Fig. 4A–D**). This binding is putatively held together by one hydrogen bond between CTC and T268 on one of the E protein chains, and by hydrophobic bonds between CTC and residues on both chains of the ZIKV E protein (**Fig. 4B**). The binding site appears to be a large pocket under the central helices on the side of the E protein that faces the viral envelope (**Fig. 4A,C**). Three of the top nine predicted binding modes (**Supplemental Fig. S1A**) indicated binding to this pocket. Notably, four out of the top nine predicted binding modes also suggested potential CTC-binding sites on the surface-exposed side of the E protein. Three of these models suggest energetically favorable binding to a negatively charged region within the cavity proximal to the central dimer interface (**Supplemental Fig. S1B**).

**Figure 4.**
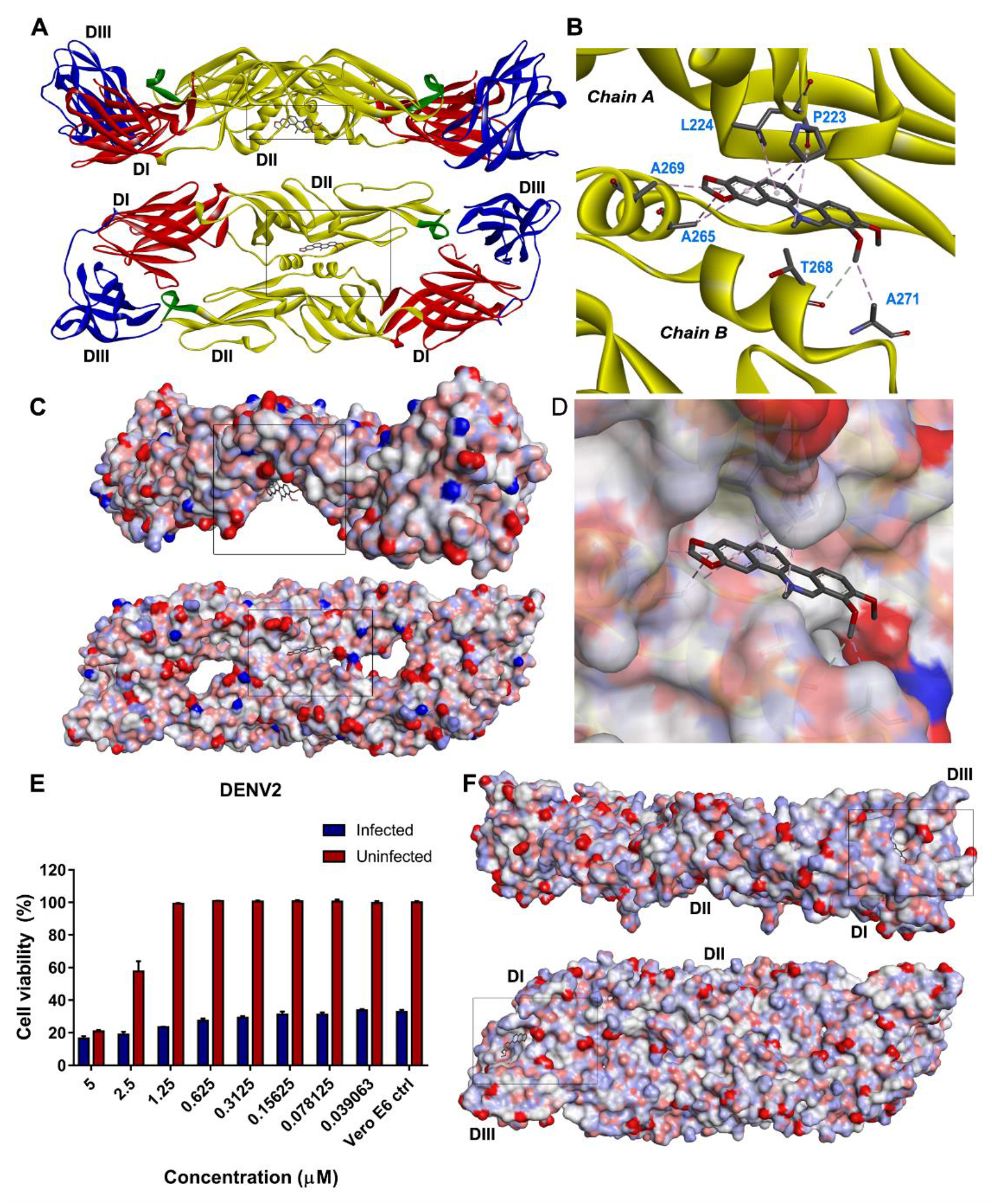
Predicted binding sites of CTC on ZIKV and dengue virus serotype 2 (DENV2) envelope (E) proteins. **(A)** Ribbon model of the CTC-binding site on the ZIKV E protein ectodomain. Top: side-view; bottom: bottom-view; DI-DIII: E protein domains 1 to 3. **(B)** Contact residues of CTC on the ZIKV E protein. **(C)** Surface model of the CTC-binding site on the ZIKV E protein in the same orientation as in **A**. Red indicates the negatively charged patches; blue indicates the positively charged patches. **(D)** Surface model of the CTC-binding pocket on the ZIKV E protein. **(E)** Effects of different concentrations of CTC on DENV2-induced (multiplicity of infection of 0.005) cytopathic effects in Vero E6 cells. Cell viability was calculated relative to the uninfected Vero E6 cells with 1% DMSO (Vero E6 ctrl). Datapoints represent the mean ± SEM; *N* = 3 per concentration. Results shown are representative of 3 independent experiments. **(F)** Predicted binding site of CTC on the surface model DENV2 E protein ectodomain. Top: side-view; bottom: bottom-view. Red indicates the negatively charged patches; blue indicates the positively charged patches.

Because CTC can inhibit WNV and ZIKV infection *in vitro*, we next investigated whether CTC inhibits DENV2 infection to determine its potential as a pan-flaviviral inactivator. We found that CTC treatment did not inhibit DENV2-induced CPE in Vero E6 cells (**Fig. 4E**). Prediction of a potential CTC-binding site on the DENV2 E protein suggested binding at a pocket distal to the central dimer interface between the DI and DIII domains of DENV2 E (PDB: 1OKE)(**Fig. 4F**)(31). We also noted that the pocket underneath the central helices of the E protein was less pronounced in DENV2 than in ZIKV. Furthermore, the cavities on either side of the central helices in ZIKV E protein, which were also potential binding sites for CTC, were less prominent in the DENV2 E protein structure (30). Only a single-chain structure of the WNV E protein was available at the time of writing, so we could not identify the potential binding sites of CTC on the dimeric prefusion form of the WNV E protein. However, our *in silico* docking analyses suggest that although CTC inhibits WNV and ZIKV infection, CTC may not be able to inhibit all flaviviral infections owing to structural differences among the flaviviral E proteins.

## Discussion

Despite the currently low number of reported ZIKV cases, ZIKV remains a threat to public health. Therapeutic agents would reduce the morbidity associated with ZIKV infection and would be helpful in controlling a future ZIKV epidemic. In this study, we show that CTC inhibits ZIKV infection in Vero and A549 cells and should be considered a potential therapeutic agent for ZIKV infection.

Further, we report that treatment with CTC renders the ZIKV particles incapable of attaching/binding to Vero and A549 cells. The effect of CTC on ZIKV is likely independent of its ability to inhibit PKC, as other PKC inhibitors did not protect Vero cells from ZIKV infection. Instead, CTC may bind to the ZIKV E protein, which prevents the interaction of the E protein with host attachment factors and/or receptors.

CTC is a cell-permeable benzo[c]phenanthridine alkaloid from plants and is considered a broad-spectrum PKC inhibitor. Previous studies have reported the potential applicability of CTC for the inhibition of RVFV and WNV infection, and these studies suggest that the inhibitory effects of CTC on these viruses are primarily due to its ability to inhibit PKC. A recently published study has also reported the inhibitory activity of CTC on ZIKV, with an EC_50_ of 779 nM on ZIKV-induced CPE in Vero cells; however, the authors did not further explore the mechanisms underlying the effects of CTC (32). The RVFV study demonstrated the inhibitory effects of CTC and other broad-spectrum PKC inhibitors (calphostin C and rottlerin) on RVFV infection in mammalian cells, clearly indicating that the RVFV replication cycle involves PKC, especially PKCε (21). Pan-PKC inhibitors, specifically CTC and calphostin C, but not inhibitors of specific PKC isoforms, were also able to inhibit WNV infection in Vero cells, suggesting that WNV also requires PKC for infection. In contrast, as we have shown here, neither rottlerin nor calphostin C reduced ZIKV-induced CPE in Vero cells. Other PKC inhibitors also did not protect Vero cells from ZIKV-induced CPE. While safingol showed inhibitory effects on ZIKV infection, the effects of safingol were modest compared to the effects of CTC. Additionally, safingol has non-PKC targets; safingol is an analog of sphingosine and can therefore act as an inhibitor of sphingosine kinases (SphK1 and SphK2), which are involved in the ceramide/sphingomyelin pathway. Growing evidence shows that flaviviruses modulate the ceramide/sphingomyelin pathway during infection (33). Thus, the inhibitory effects of safingol on ZIKV-induced CPE may be partially attributed to its effects on this pathway and may explain why safingol showed inhibitory effects on ZIKV infection while the other PKC inhibitors did not. Taken together, our results suggest that ZIKV and WNV have different requirements for PKC, where WNV may be more dependent than ZIKV on PKC for infection.

The failure of other PKC inhibitors to inhibit ZIKV infection in Vero cells led us to speculate on a PKC-independent target for CTC. Further investigation revealed that CTC inhibits ZIKV attachment/binding rather than internalization in Vero cells. Pretreatment of cell-free ZIKV particles with CTC reduced ZIKV attachment/binding to Vero and A549 cells, verifying our hypothesis that PKC-independent and host-independent mechanisms are involved in the inhibitory effects of CTC on ZIKV infectivity. Prediction of the interaction between CTC and the prefusion form of the ZIKV E protein suggests that CTC binds a pocket underneath the central helices of the ZIKV E protein. The prefusion ectodomain of the ZIKV E protein lies flat against the viral envelope, and our top predicted binding model suggests that CTC binds to a pocket on the side facing the viral envelope. While there may be accessibility constraints based on the location of this pocket, the putative binding pocket for CTC appears deep; furthermore, the prefusion conformation of the flavivirus E protein undergoes “molecular breathing,” which suggests a highly dynamic state that promotes contact with the environment (34,35). We speculate that the size of this pocket, the dynamic state of the E protein, and the cavities on either side of the central dimer interface allow CTC to access the pocket underneath the helices despite potential structural constraints. Furthermore, docking prediction suggested another binding site on the cavity proximal to the central dimer interface. These cavities are on the surface-exposed side of the ZIKV E protein, and CTC may bind to these more accessible cavities instead. Whether CTC binds to the envelope-facing or the surface-exposed side of the E protein, binding of CTC to the central dimer region of the ZIKV E protein may stabilize the E protein to a more rigid conformation, making it unable to interact with attachment factors and binding receptors on the host cell surface, ultimately reducing the infectivity of ZIKV particles.

In contrast, the predicted binding site of CTC on the DENV2 E protein is on the ends of the prefusion conformation of the E protein. Given the tight end-to-end formation of the flaviviral E proteins at the prefusion state, this binding site may be inaccessible to CTC in the context of the intact virion, which may explain the differences between the effects of CTC on ZIKV and DENV2 infection *in vitro*. Whether CTC similarly binds the WNV E protein and whether this mechanism contributes to the inhibitory effects of CTC on WNV infection should be investigated. The effects of CTC on other flaviviruses should also be evaluated to allow us to determine the range of targets that CTC inhibits, at least within this family of viruses.

The host-independent activity of CTC against ZIKV and docking prediction with the ZIKV E protein indicates that the CTC structure influences its inhibitory effects on ZIKV. Thus, identification of the CTC pharmacophore that binds the ZIKV E protein will allow modification of CTC for optimal pharmacokinetic and pharmacological properties. Furthermore, compounds with structures or properties similar to CTC can be screened for their ability to reduce ZIKV infectivity. In line with this, berberine, an alkaloid that is structurally related to CTC, has been reported to exhibit virucidal effects on ZIKV (36).

Although targeting host-dependent mechanisms of infection may reduce the likelihood of the development of viral resistance to drugs (37), they may result in adverse effects, especially if the drugs affect several biological pathways (38). Therefore, drugs that directly target viruses have the advantage of minimizing drug-related side effects. CTC is still at the pre-clinical stage of development, primarily as an anticancer agent. Little information on its pharmacokinetics and safety is available. However, a study has shown that long-term treatment with high doses of CTC (≥5.6 mg/kg) caused lung damage in a rat model (39). By contrast, no deaths were observed among mice treated with CTC at 2.5–5.0 mg/kg/day for three days (40). *In vivo* studies will have to be performed to determine whether safe doses of CTC will protect animal models, especially fetal development models, from ZIKV infection.

In this study, we demonstrated the anti-ZIKV activity of CTC, a known PKC inhibitor, *in vitro*. However, other PKC inhibitors failed to protect Vero cells from ZIKV-induced CPE in Vero cells, suggesting that PKC-independent mechanisms are responsible for the inhibitory effects of CTC on ZIKV. Treatment of cell-free ZIKV particles with CTC reduced ZIKV attachment/binding to Vero and A549 cells, which supports our hypothesis that PKC- and cell-independent mechanisms are at play. Specifically, CTC may bind to the ZIKV E protein and stabilize the prefusion state of the E protein, thereby inhibiting the interaction of the E protein with attachment and binding factors on the host cell surface. This potential mechanism has not been previously reported, as most studies on the antiviral effects of CTC suggest PKC-dependent mechanisms. Binding assays should be performed to determine whether CTC occludes ZIKV E protein attachment/binding to host factors or if other mechanisms cause the inactivation of ZIKV particles by CTC. Furthermore, we focused primarily on the effects of CTC on ZIKV entry using *in vitro* models and did not determine the effects of CTC on post-fusion stages of infection. Whether CTC targets later stages in the ZIKV replication cycle and whether these effects are PKC-dependent should be verified in future studies to help us understand the extent of the inhibitory effects of CTC on ZIKV infection. Overall, our results indicate that CTC reduces the infectivity of ZIKV particles *in vitro*. These findings support further exploration of CTC as a potential therapeutic agent for ZIKV infection.

## Materials and Methods

### Cells and viruses

African green monkey (Vero; ATCC CCL-81), A549 (ATCC CCL-185), and Vero E6 (ATCC CRL-1586) cells were obtained from the American Type Culture Collection (ATCC; Manassas, VA). These cells were propagated in growing medium consisting of minimal Eagle’s medium (MEM; Gibco, Carlsbad, CA) supplemented with 10% fetal bovine serum (FBS) (Gibco), antimycotic antibiotics (Gibco), and ʟ-glutamine (Gibco). ZIKV (ATCC VR-1838) was propagated in Vero cells using MEM supplemented with 0.3% bovine serum albumin (BSA), antimycotic antibiotics, and ʟ-glutamine (MEM + BSA) at 37 °C in a humidified atmosphere with 5% CO_2_. The ZIKV particles were harvested at 5 days post-infection (dpi), and culture supernatants were stored at –80 °C until use. MEM supplemented with 1% FBS (MEM + 1% FBS), antimycotic antibiotics, MEM vitamins, and ʟ-glutamine was used to infect A549 cells. DENV2 (ATCC VR-1584) was propagated in Vero E6 cells in MEM + 1% FBS for 5 d at 37 °C in a humidified atmosphere with 5% CO_2_. The DENV2 particles were harvested and titrated in Vero E6 cells via plaque assay, and culture supernatants were stored at –80 °C until use.

### Reagents

CTC was purchased in the form of chelerythrine chloride from Sigma-Aldrich (C2932; St. Louis, MO) and reconstituted to 5 mM in DMSO. Stock solutions (5 mM and 500 μM) of CTC in DMSO were stored in aliquots at −20 °C. Bisindolylmaleimide I (Cat. No. 203291), calphostin C (208725), Gö 6976 (Cat. No. 365250), Gö 6983 (Cat. No. G1918), and rottlerin (Cat. No. 557370) were purchased from Sigma-Aldrich. Safingol (Cat. No. 559300) was purchased from Merck-Millipore (Burlington, MA). Bisindolylmaleimide I, calphostin C, Gö 6976, and Gö 6983 were reconstituted to 1 mM in DMSO, safingol was reconstituted to 10 mM in DMSO, and bryostatin 1 was reconstituted to 100 μM in DMSO. All PKC inhibitors were stored in aliquots at −20 °C. CQ in the form of chloroquine diphosphate salt (Cat. No. C6628; Sigma-Aldrich) was dissolved to 100 mM in distilled deionized water and stored at 4 °C. All chemicals were diluted to working concentrations in MEM + BSA or MEM + 1% FBS on the day of use. The cytotoxicity assay kit EZ-CYTOX was purchased from DoGenBio Co., Ltd. (Seoul, South Korea).

### Dose-dependent effects of PKC inhibitors on ZIKV-induced CPE in Vero cells

Vero cells (1.2 × 10^4^ cells/well) were grown overnight in 96-well plates. The cells were then washed with phosphate-buffered saline (PBS). PKC inhibitors were diluted to the indicated concentrations in MEM + BSA. PKC inhibitors were added to Vero cells in the presence of ZIKV at an MOI of 0.02 to a total volume of 100 μL/well (*N =* 3 per concentration), with 1% DMSO for all treatments. The cells were incubated at 37 °C with 5% CO_2_ for 5 d. After incubation, EZ-CYTOX (10 µL/well) was added, cells were incubated at 37 °C with 5% CO_2_, and absorbance at 450 nm was measured after 3– 4 h. Percent cell viability was calculated relative to uninfected Vero cells treated with 1% DMSO. To obtain the EC_50_, percent cell viability values were normalized relative to the uninfected Vero control with 1% DMSO (maximum) and the infected Vero control with 1% DMSO (minimum) per assay using GraphPad Prism version 7.0 (GraphPad Software, San Diego, CA). The EC_50_ of CTC against ZIKV-induced CPE was calculated based on three independent assays using the nonlinear regression-variable slope function in GraphPad Prism version 7.0.

### Cytotoxic effects of PKC inhibitors in Vero cells

Vero cells (1.2 × 10^4^ cells/well) were grown overnight in 96-well plates. The cells were washed with PBS. The PKC inhibitors were diluted in MEM + BSA to the indicated concentrations and added to Vero cells to a final volume of 100 μL/well with 1% DMSO (*N =* 3 per concentration). The cells were incubated at 37 °C with 5% CO_2_ for 5 d. After incubation, EZ-CYTOX (10 µL/well) was added. Cells were incubated at 37 °C with 5% CO_2_, and the absorbance at 450 nm was read after 3–4 h. Percent cell viability was calculated relative to uninfected Vero cells treated with 1% DMSO. For CC_50_ calculation, cell viabilities were normalized relative to the Vero control treated with 1% DMSO (maximum), and to the minimum cell viability per assay using GraphPad Prism version 7.0. The CC_50_ of CTC in Vero cells was calculated based on three independent assays using the nonlinear regression-variable slope function of GraphPad Prism version 7.0.

### ZIKV growth kinetics assay in Vero cells

Vero cells (2.5 × 10^5^ cells/well) were prepared in 6-well plates and incubated overnight at 37 °C with 5% CO_2_. The cells were washed once with PBS. Then, ZIKV (MOI of 0.01) was added to the Vero cells in the presence of 1.45 μM CTC, 25 μM CQ, or 0.3% DMSO to a total volume of 3 mL per well (*N =* 3 per treatment) in MEM + BSA. The cells were incubated for 96 h at 37 °C, with 5% CO_2_.

Supernatants (250 μL) were collected at 6, 24, 48, 72, and 96 hpi. At 96 hpi, CTC- and DMSO-treated cells (with and without ZIKV) were observed under a light microscope to determine the effects of CTC treatment on cell morphology. Supernatants were stored at −80 °C prior to plaque assay.

### Determination of ZIKV titer via plaque assay

Vero cells (1.5 × 10^5^ cells/well in 12-well plates or 3.5 × 10^5^ cells/well in 6-well plates) were incubated at 37 °C with 5% CO_2_ for two days (95% confluence). The cells were washed with PBS, and then inoculated with 250 μL (12-well plate) or 1 mL (6-well plate) of 10-fold serially diluted ZIKV infection supernatants (*N =* 3) in MEM + BSA for 2 h at 37 °C, with 5% CO_2_. The infection supernatant was removed, and the cells were overlaid with MEM + BSA with bacteriological agar (0.9%). Cultures were incubated at 37 °C with 5% CO_2_ for 96 h, after which the agar overlay was removed. The plaques were stained and fixed in 0.1% crystal violet and 10% formaldehyde. The mean titer (PFU/mL) for each treatment was calculated. The limit of detection for this assay was 10 PFU/mL.

### Dose-dependent inhibition of ZIKV production in A549 cells

A549 cells (2.5 × 10^5^ cells/well) were prepared in 24-well plates and incubated overnight at 37 °C with 5% CO_2_. The cells were then washed once with PBS. Then, ZIKV (MOI of 0.01) was added to the cells in the presence of different concentrations of CTC or with 1% DMSO diluted in MEM + 1% FBS (total of 1 mL/well; *N =* 3/concentration). The cells were incubated at 37 °C for 72 h. A549 culture supernatants were collected and subjected to plaque assay in Vero cells to determine ZIKV titers.

### Time-of-drug-addition assay in Vero cells

The time-of-drug-addition assay was based on a previous study with modifications^16^. Vero cells were prepared overnight in 24-well plates (1.5 × 10^5^ cells/well). The cells were washed once with PBS and infected with ZIKV (MOI of 10.0) for 1 h at 4 °C. The cells were washed twice with ice-cold PBS to remove unbound virus particles. MEM + BSA (500 µL/well) was added to all wells following adsorption. The drugs diluted in MEM + BSA (1.45 μM CTC or 25 μM CQ; 500 µL/well) containing 0.3% DMSO were added at different time points post-infection (0, 0.5, 3, 6, 12, and 24 hpi; *N =* 3 per time point). The untreated, infected control was prepared by the addition of 0.3% DMSO (1 mL/well) in MEM + BSA at time 0 (*N =* 3). Culture supernatants were collected at 30 hpi and stored at –80 °C prior to plaque assay.

### Attachment/binding inhibition assay in Vero cells

Vero cell cultures were prepared in 6-well plates (3.5 × 10^5^ cells/well) two days before the assay (until 95% confluence). The cells were then washed once with PBS. The cells were infected with 400 PFU per well of ZIKV in the presence of 1.45 μM CTC, 2.9 μM CTC, 25 μM CQ, or 1% DMSO in MEM + BSA (1 mL total volume) for 2 h at 4 °C **(Fig. 3*B*)**. The ZIKV + drug mixture was removed, and the cells were washed twice with ice-cold PBS to remove unbound virus particles. The plaque overlay was added (3 mL/well), and the cells were incubated at 37 °C for four days prior to staining.

### Attachment/binding inhibition assay in A549 cells

Overnight cultures of A549 cells (2.5 × 10^5^ cells/well) were prepared in 24-well plates. The cells were then washed once with PBS. Different concentrations of CTC (500 μL/well; *N =* 3 per concentration) or 1% DMSO in MEM + 1% FBS and 500 μL of ZIKV (MOI of 1.0) in MEM + 1% FBS were added to the cells. The virus particles were allowed to adsorb to A549 cells for 2 h at 4 °C, and the cells were washed twice with ice-cold PBS to remove unbound virus particles. The infected cells were incubated at 37 °C, and culture supernatants were collected after 24 h. Culture supernatants were stored in aliquots at –80 °C. Production of ZIKV in A549 cells was determined by plaque assay in Vero cells.

### Internalization inhibition assay in Vero cells

Vero cell cultures were prepared in 6-well plates (3.5 × 10^5^ cells/well) two days before the assay (until 95% confluence). The cells were washed once with PBS and then infected with ZIKV (400 PFU in 1 mL MEM + BSA per well) at 4 °C for 2 h **(Fig. 3*B*)**. The inoculum was removed, and the cells were washed twice with ice-cold PBS. The cells were treated with 1.45 μM CTC, 2.9 μM CTC, 25 μM CQ, or 1% DMSO (1 mL in MEM + BSA) for 2 h at 37 °C. The drug was removed. The cells were washed once with PBS, treated with 300 μL citric acid buffer (40 mM citric acid, 10 mM KCl, 135 mM NaCl; pH 3.0) for 30 s, and then washed again with PBS. The plaque overlay was added (3 mL/well), and the cells were incubated at 37 °C for four days prior to staining.

### Vero cell pretreatment assay

Vero cell cultures were prepared in 6-well plates (3.5 × 10^5^ cells/well) two days before the assay (until 95% confluence). The cells were washed once with PBS, and then treated with 1.45 μM CTC, 2.9 μM CTC, 25 μM CQ, or 1% DMSO (1 mL in MEM + BSA) for 2 h at 37 °C. The drugs were removed, and the cells were washed twice with PBS. The cells were then infected with ZIKV (400 PFU/mL/well in MEM + BSA) for 2 h at 4 °C. The inoculum was removed, and the cells were washed twice with ice-cold PBS to remove unbound virus particles. The plaque overlay (3 mL/well) was added, and the cells were incubated for four days at 37 °C prior to staining.

### Virus pretreatment assay for Vero cell infection

The virus inactivation assay was based on the protocol of Tai *et al. (*41). Vero cell cultures were prepared in 6-well plates (3.5 × 10^5^ cells/well) two days before the assay (until 95% confluence). ZIKV particles (3.75 × 10^4^ PFU) in 250 μL MEM + BSA were mixed with 250 μL of CTC, CQ, or DMSO in MEM + BSA to final concentrations of 1.45 μM CTC, 25 μM CQ, or 1% DMSO. The particles were treated for 0, 2, or 4 h at room temperature (*N =* 3 per time point) (**Fig. 3*B***). All time points had corresponding 1% DMSO controls (*N =*3 per timepoint) to account for loss of viral particles over the incubation period. Immediately before infection, the compound + ZIKV mixture was serially diluted to obtain a final dilution of 900 PFU in 1.2 mL of MEM + BSA and 1:100 dilution (subtherapeutic concentration) of the drug.

The cells were washed once with PBS before infection. Then, 1 mL of the diluted ZIKV (750 PFU/mL) + drug mixture was added to Vero cells. The cells were incubated at 4 °C for 2 h to facilitate binding. The cells were washed twice with PBS to remove unbound virus particles, and the plaque overlay (3 mL/well) was added. The cells were incubated at 37 °C for 4 d prior to staining.

### Virus pretreatment assay for A549 cell infection

Overnight cultures of A549 cells (2.5 × 10^5^ cells/well) were prepared in 24-well plates and washed once with PBS before infection. On the day of the assay, ZIKV particles (8.0 × 10^4^ PFU) were treated with 2.5 μM CTC in a total volume of 400 μL in MEM + BSA at room temperature for 0, 2, or 4 h before infection (*N =* 3/time point). Immediately before infection, the CTC + ZIKV mixtures were diluted 1:80 in MEM + 1% BSA to reduce CTC to subtherapeutic levels. The diluted ZIKV + CTC mixtures were used to infect A549 cells (1,250 PFU ZIKV/well) for 2 h at 4 °C. The cells were washed twice with ice-cold PBS to remove unbound virus particles and were incubated at 37 °C for 24 h. ZIKV particles incubated with 1% DMSO were likewise prepared for all time points (*N =* 3/time point) to account for the loss of virus in cell-free conditions at room temperature. At 24 hpi, culture supernatants were collected and subjected to plaque assay in Vero cells to determine ZIKV titers.

### CTC cytotoxicity in Vero E6 cells

Vero E6 cells (2.5 × 10^4^ cells/well) were grown overnight in 96-well plates. The cells were washed with phosphate-buffered saline (PBS). CTC was diluted in MEM + 1% FBS to the indicated concentrations and added to Vero E6 cells to a final volume of 100 μL/well with 1% DMSO (*N =* 3 per concentration). The cells were incubated at 37 °C with 5% CO_2_ for 5 d. After incubation, EZ-CYTOX (10 µL/well) was added. Cells were incubated at 37 °C with 5% CO_2_, and the absorbance at 450 nm was read after 3–4 h. Percent cell viability was calculated relative to uninfected Vero E6 cells treated with 1% DMSO.

### CTC activity against DENV2 in Vero E6 cells

Vero E6 cells (2.5 × 10^4^ cells/well) were grown overnight in 96-well plates. The cells were then washed with PBS. CTC was diluted to the indicated concentrations in MEM + 1% FBS. The drug was added to Vero cells in the presence of DENV2 at an MOI of 0.005 to a total volume of 100 μL/well (*N =* 3 per concentration), with 1% DMSO for all treatments. The cells were incubated at 37 °C with 5% CO_2_ for 5 d. After incubation, EZ-CYTOX (10 µL/well) was added, the cells were incubated at 37 °C with 5% CO_2_, and absorbance at 450 nm was measured after 3–4 h. Percent cell viability was calculated relative to uninfected Vero E6 cells treated with 1% DMSO.

### Docking prediction

AutoDock Vina included in the PyRx v.8.0 (42) package was used to predict the docking sites of CTC (PubChem CID: 2703) on the prefusion conformation of ZIKV (PDB: 5JHM)(30) and DENV2 E proteins (PDB: 1OKE)(31). Default parameters were used, and the top nine binding modes were selected. Discovery Studio Visualizer v.21.1 (Biovia Dassault Systèmes, Vélizy-Villacoublay, France) was used to view the docking results.

### Statistical analyses

Student’s *t*-test was used to determine significant differences between pairs of groups (*P* < 0.05). One-way analysis of variance (ANOVA), followed by Tukey’s multiple comparison test, was used to test for significant differences among groups (*P* < 0.05). Statistical analyses were performed using GraphPad Prism v.7.0.

## Acknowledgments

This work was supported by the National Research Foundation of Korea (NRF) grant funded by the Korea government (MSIT) (No. 2022R1A2C1011553), the Korea University Grant (K2112821 and K2207131) and ALSAC. Figure 3B was created with BioRender.com.

