## Supplement Figure S1 for "Chelerythrine as an anti-Zika virus agent: therapeutic potential and mode of action"

**Supplemental Figure**

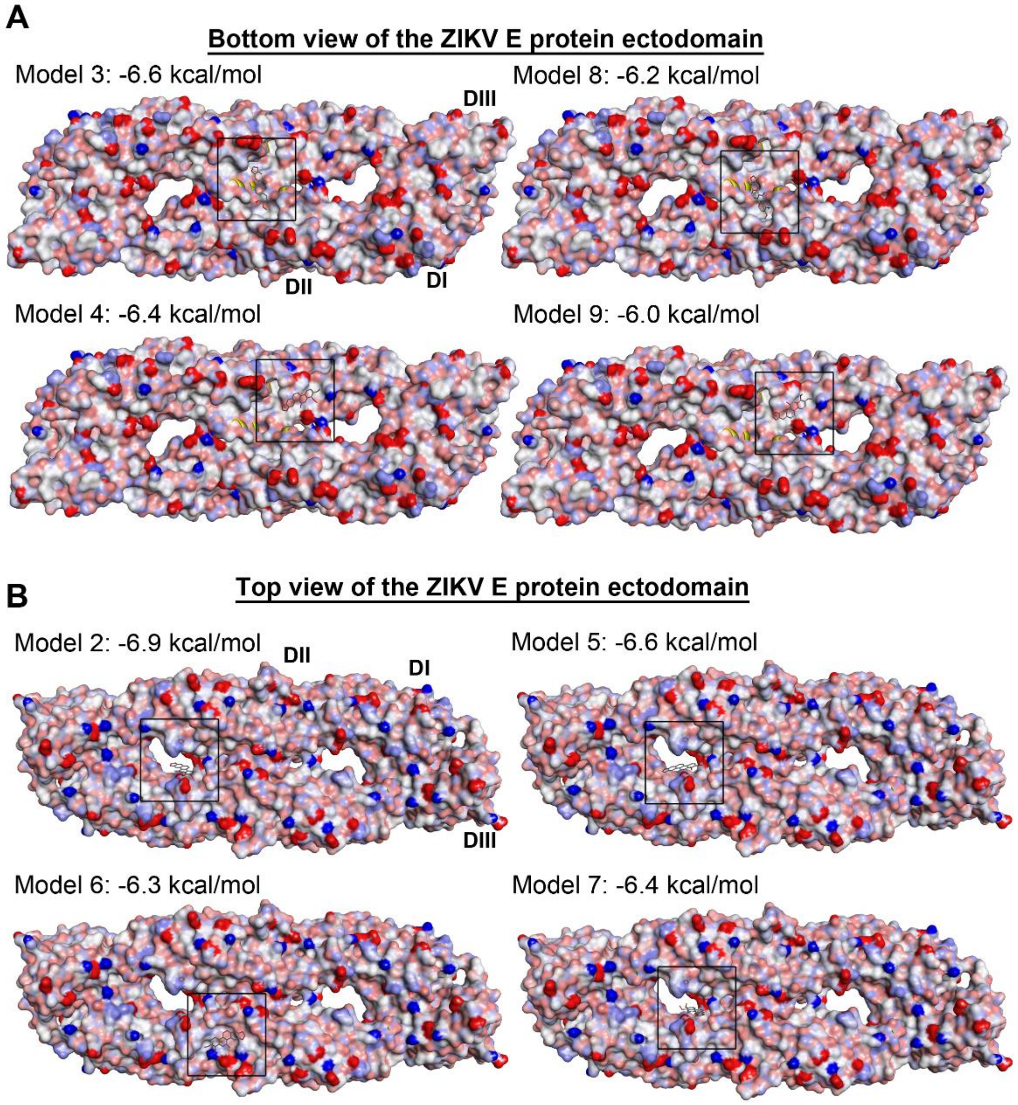


**Fig. S1. Other predicted binding models of chelerythrine (CTC) on the Zika virus (ZIKV) envelope (E) protein. (A)** Models 3, 8, 4, 9 as viewed from the bottom (envelope-facing side) of the E protein. Models 3 and 8 suggest CTC binding (boxed) to the underside of the central helices of the E protein. Models 4 and 9 indicate binding to the beta sheet region (boxed) of the ZIKV E protein domain II (DII). **(B)** Models 2, 5, 6, and 7 as viewed from the top (surface-exposed side) of the E protein ectodomain. Models 2, 5, and 7 show CTC binding (boxed) on a negative patch on a cavity proximal to the central dimer interface in the E protein. Domains I, II, and III (DI–DIII) belonging to a single chain of the ZIKV E protein are labelled. Calculated binding affinities (kcal/mol) are indicated. Red regions on the surface model indicate negatively charged patches; blue regions indicate positively charged patches. Binding sites were predicted using PyRx version 7 and visualized with Discovery Studio Visualizer version 21.
